# Rob and MarA alter susceptibility of *Escherichia coli* to antibiotics in presence of salicylate

**DOI:** 10.1101/141671

**Authors:** Kirti Jain, Supreet Saini

## Abstract

When exposed to stress, bacterial cells launch a diverse response to enhance their chances of survival. This response involves modulation of expression of a large number of proteins which help the cell counter stress. This modulation is facilitated by several transcription factors in bacteria and in *E. coli* three homologous regulators, MarA, Sox, and Rob are known to launch a coordinated response to combat various stress environments. MarA and SoxS are known to control multiple antibiotic resistance and superoxide regulon respectively. Rob has been observed to control similar downstream targets as MarA and SoxS. However, physiological relevance of Rob in not understood. We show that Rob along with MarA, in presence of inducer salicylate, can help cell survive in presence of lethal concentration of wide range of antibiotics.

## Introduction

Transcriptional regulation of cellular targets like efflux transporters or outer membrane porins help a bacterial cell combat stress [1–4]. In *E. coli*, three homologous transcriptional regulators – MarA (encoded by *multiple antibiotic resistance marRAB* operon), SoxS (encoded by superoxide stress *soxSR* regulon), and Rob (right origin binding protein), together known as *mar/sox/rob* regulon, are known to regulate these processes [1,5–11]. For e.g., the TolC efflux pump is known to be upregulated by MarA, SoxS, and Rob [4]. Production of porin protein OmpF is regulated by small RNA *micF* [1], which is regulated by MarA, Sox, and Rob. Many of the cellular enzymes mentioned like Zwf, DeoB, SodA, etc. are also known to be controlled by either of MarA, SoxS, or Rob [12–14]. MarA and SoxS have been reported to play a role in conferring antibiotic resistance and combating superoxide stress [5,15–21]. Rob is known to act on an overlapping set of targets as MarA and SoxS [7,22–24], however, its role in regulating cellular physiology is not clearly understood [25,26].

The *marRAB* operon is known to be induced in presence of compounds such as salicylate, phenolic compounds [4,5,8,27–29]. *marRAB* system encodes for a transcriptional activator MarA, repressor MarR, and protein of unknown function MarB [8,15–17,29,30]. MarR repressor, in absence of inducers, remain bound to Pmar promoter and represses expression from the *Pmar* promoter [15–17]. In presence of inducers, MarR preferentially binds to the inducer molecule, hence relieving the repression of the *Pmar* promoter [15–17,23]. Thereafter, MarA binds to the Pmar and target genes promoters and activates transcription [1,4–6,23,27,28,31].

Rob, (right origin binding protein), was first discovered to bind to a site close to DnaA binding site to the DNA, [25]. However, its precise role in controlling DNA replication is not known [25]. Crystal structure of Rob further revealed structural homology with transcriptional regulators MarA and Sox and has been classified as AraC/XylS type transcriptional regulator family [7,12,13,22–24,32]. However, unlike MarA and SoxS, transcription from the *rob* promoter is constitutive (the protein in its uninduced state is present in the cell as agglomerate) and the protein amounts are regulated post translationally, in presence of cognate inducer bile slats [12,13,33,34]. Salicylate is also known to control gene expression via Rob, however, the molecular mechanism for the same is not understood [35,36].

Previous work from our lab demonstrated that in presence of salicylate, MarA and Rob together control downstream targets like InaA by forming a Feed Forward Loop [35]. MarA protein is important in altering MIC value of a large number of antibiotics in gram-negative bacteria. However, physiological significance of Rob in this context is not well understood. In this work, we show that Rob plays a similar role as MarA in altering the MIC of antibiotics in bacteria in presence of the canonical inducer salicylate. Since salicylate is the active molecule of aspirin (acetyl salicylic acid), a clinically important molecule, it is important to understand the molecular mechanism of this action and its further impact on antibiotic resistance.

## Methods

### Growth Kinetics

Cells were grown in LB media overnight with shaking at 37°C. The overnight cultures were diluted 1:250 in fresh media. Cells were thereafter allowed to grow till an OD 0.2 upon attaining which and respective inducer was added. Growth dynamics was captured for – wild type, Δ*marA*, and Δ*rob* in induced and uninduced conditions. A range of inducer, salicylate (Sigma Aldrich) concentration - 1mM, 5mM, and 10mM) was taken to capture sub-lethal effect by inducers. Experiments were performed for a range of inducer concentrations to capture their precise physiological effect at different concentrations.

### Minimal Inhibitory Concentration (MIC) determination

Broth microdilution method was used to determine MIC of an antibiotic. Antibiotic stocks and the concentration range used were prepared as per the guidelines given by National Committee for Clinical Laboratory Standards, NCCLS [37]. Antibiotic concentration range was taken one fold higher than the required range to count for dilution because of inoculum addition. The inoculum was prepared using an overnight culture (LB media) by diluting to have 10^6^ cells per ml. A sterile 96 well plate was taken and 100μL of inoculum was added to the well containing 100μL of fresh media with appropriate antibiotic concentration. Thus, the final cell density for the experiment was kept at 5 × 10^5^ cells per ml. Inoculum density was cross checked every time by counting the colony forming unit (CFU) on LB agar plate. The plate was sealed using breathe easy membrane (Sigma Aldrich) and incubated at 37°C for 20 hours. The concentration of the lowest antibiotic concentration well showing no visible growth was interpreted as the MIC for a particular antibiotic.

### Live-Dead Cell Assay using Propidium Iodide (PI)

Cells were exposed to antibiotics in presence and absence of salicylate to track the percentage of dead cells using propidium iodide assay (Sigma Aldrich). Propidium iodide is known to rapidly penetrate cells with compromised membrane or dead cells. Once in touch with DNA it fluoresces at 617nm. The reason for choosing this dye is its minimal interference with inducers or antibiotics used in this study.

An overnight culture of wild type and mutants was sub-cultured in 1:250 dilution. After 1.5 hours of growth at 37°C with shaking, OD 600 was monitored every 10 minutes using OD density meter. When the OD reached ~0.4, the cells were exposed to salicylate at a concentration of 7.5mM. After an hour of salicylate exposure (OD ≤ 0.65), cells were exposed to varying concentration of antibiotics. The antibiotic range was decided as depending on the MIC values for each antibiotic. For the purpose of this study, the antibiotic range was chosen to consider both sub-lethal and lethal concentrations of antibiotics; i.e., both < MIC and > MIC concentration. The percentage of dead cells was estimated after three hours of antibiotic exposure, roughly in mid-exponential phase. Samples were collected and stored in PBS. Propidium iodide was used at a concentration 10μg/ml. Samples were immediately observed at single-cell resolution using a Flow Cytometry (Millipore Guava System and BD FACS Aria SORP). 50,000 events were recorded and FSC/SSC were plotted. Care was taken to not expose samples to light after adding Propidium Iodide. All single-cell expression experiments were repeated thrice on different days and found to show similar trend.

## Results

### *mar* and *rob* systems are associated with a phenotype in *E. coli*

To characterize the physiological role of MarA and Rob, growth dynamics were performed for wild type and mutants (in presence and absence of inducers) and growth dynamics compared. These downstream targets control several aspects of cellular physiology and consequently have an effect on growth dynamics of the bacterium. Previous reports suggest that MarA and Rob control a number of common downstream cellular targets [4,6,22,23,38–40]. Hence, the changes in growth phenotype arising due to absence of these systems were explored by measuring the growth defect associated with deletion of *marA* or *rob*.

Kinetic experiments were carried out to see the difference in growth dynamics in regulatory mutants (Δ*rob* and Δ*marA*) in presence and absence of salicylate, and compared to that of wild type. Our results show that, compared to wild-type *E. coli*, growth of mutants Δ*marA* and Δ*rob* is inhibited in presence of sub-inhibitory concentrations of salicylate. This effect is especially pronounced at lower concentrations of salicylate. Defect in growth is calculated as percentage growth difference in induced cells as compared to uninduced conditions (Figure 1). At low concentration of salicylate (1mM), the wild-type cells show growth defect of around 10% as compared to 15–20% for mutants Δ*rob* and Δ*marA* (Figure 1B and 1C). For higher concentrations of salicylate (5mM and 10mM), wild type and mutants show comparable growth defects. Since *mar/sox/rob* systems are thought to have evolved to combat sub-lethal stress [28,29,41], it is not surprising that absence of either regulator leads to growth defect when exposed to low inducer concentration (1mM) as compared to wild-type.

**Figure 1.**
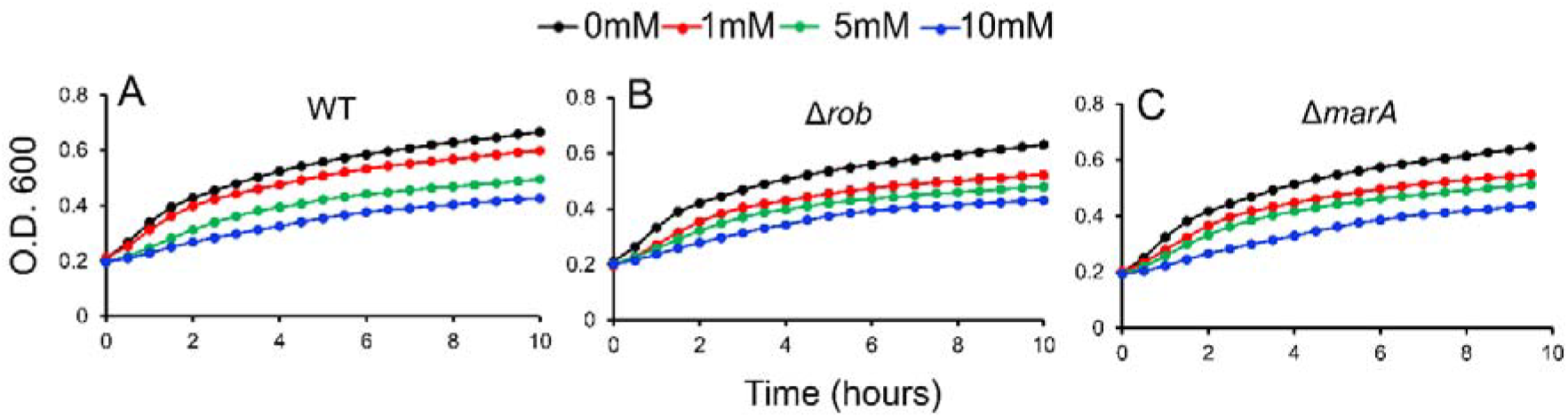
Growth defect in wild-type ***E. coli*** and mutants in presence of varying concentrations of salicylate. (**A**) Wild type (**B**) *Δrob*, and (**C**) Δ*marA*. Salicylate concentration used: 0mM, 1mM, 5mM, and 10mM.

### Salicylate alters Minimal Inhibitory Concentration (MIC) of antibiotics via both, MarA and Rob

Another approach to analyze the phenotype of *mar*/*rob* is to study their role in changing the Minimal Inhibitory Concentration (MIC) of antibiotics. MIC is defined as the lowest concentration of the antibiotic at (and beyond) which the organism cannot grow. One of the first published reports regarding role of salicylate and other similar compounds in altering MIC of known antibiotics was by Rosner in 1985 [41]. Since then a number of reports focus on the role of salicylate and acetyl salicylic acid in reducing susceptibility of the bacterium to antibiotics like fluroquinolone, carbenecillin, carbapenem, etc. [42,43]. However, there is no systematic work establishing role of inducers of Rob in changing MIC.

To understand the role of salicylate in altering MICs, we selected three antibiotics, commonly used in clinical practice against gram-negative bacteria. Carbenecillin is a β-lactam antibiotic having bactericidal activity, generally used against gram-negative infections. It acts by inhibiting cell wall synthesis process by interfering with the final transpeptidation step [44–46]. Cefotaxime is a broad spectrum antibiotic belonging to cephalosporin (third generation β-lactam group) active against both gram-positive and gram-negative bacteria. The mechanism of action of cefotaxime is similar to that of carbenecillin [44,45,47]. Ciprofloxacin is a fluroquinolone having bactericidal activity against gram-negative bacteria, and acts by inhibiting the process of DNA synthesis by hindering topoisomerase activity [48–51]. In this section, we focus on understanding the fold changes in MICs of antibiotics Carbenecillin, Cefotaxime, and Ciprofloxacin when both wild-type and regulatory mutant cells are grown in presence of inducers of *mar*/*rob* systems. We used salicylate as inducer.

Broth microdilution method was used to determine the MIC. Antibiotic stocks and the concentration range used were prepared as per the guidelines given by National Committee for Clinical Laboratory Standards, NCCLS [37].

To assess the effect of inducer salicylate, using MIC determination method, we first determined the sensitivity of *E. coli* to salicylate. MIC of salicylate was determined to be 4mg/ml which corresponds to a concentration of 25mM. We found that similar values of MIC have been reported previously for enterotoxigenic *E. coli* strain [52]. In our work, while using as an inducer, we have used lower concentration, 7.5mM of salicylate (represents both salicylate and acetyl salicylic acid) to prevent cell damage.

We performed broth microdilution assay for carbenecillin, cefotaxime, and ciprofloxacin both in absence and presence of inducers salicylate and paraquat to determine MIC. The concentration range of antibiotic screened and MIC values obtained are given in Table 1.

**Table 1.** MIC values for wild type and double mutants cells in presence and absence of inducer salicylate.

| MIC in µg/ml |  |  |  |  |  |
| --- | --- | --- | --- | --- | --- |
| Antibiotics | Range of concentration checked in µg/ml | Wild type |  | *ΔmarA Δrob* |  |
|  |  | W/O Salicylate | W/ Salicylate | W/O Salicylate | W/ Salicylate |
| **Carbenecillin** | .25-128 | 32 | >64 | 16 | 64 |
| **Cefotaxime** | 0.004-128 | 0.064 | 0.512 | 0.032 | 0.064 |
| **Ciprofloxacin** | 0.004-128 | 0.032 | 0.512 | 0.016 | 0.128 |

For wild-type cells, MIC for carbenecillin, was determined to be 32μg/ml. For cefotaxime, MIC was determined to be 0.06μg/ml, and for ciprofloxacin 0.032μg/ml. All three MIC values were found to be in the reported range of MIC for *E. coli* ATCC 25922 strain. Next, we repeated the same exercise by determining the respective MICs in presence of 7.5mM salicylate. We observed a two-fold change in MIC of carbenecillin in presence of salicylate. For cefotaxime and ciprofloxacin there was more than four-fold change when exposed to salicylate (Table 1). This suggests that inducer (salicylate) helps the cell survive even when exposed to lethal antibiotic concentration.

Next, we were interested in determining the MIC of the three antibiotics for single and double mutants of *ΔmarA* and *Δrob*. Deleting either of MarA or Rob did not lead to significant chance in MIC. Hence, we chose double mutant for further characterization of the role of salicylate in altering MIC via both MarA and Rob. For *ΔmarA Δrob*, for carbenecillin, cefotaxime, and ciprofloxacin, MIC was found to be half of that of wild type, i.e., 16μg/ml for carbenecillin, 0.016μg/ml for ciprofloxacin, and 0.032μg/ml for cefotaxime. The deletion of these regulators was observed to lead to at least 50% decrease in MIC when observed in uninduced condition. This suggests that the regulators might also be regulated by other metabolic intermediates as reported by Chubiz and co-workers [28].

We then repeated the same exercise in presence of 7.5mM salicylate. In absence of both MarA and Rob MIC was observed to be two-four fold less as compared to wild type MIC in identical conditions. The comparison of uninduced and induced values of MIC of all three studied antibiotics in *ΔmarA Δrob*, however, reveals around two-fold higher MIC in salicylate induced condition as compared to uninduced condition. Salicylate is known to act via MarA and Rob. The change in MIC in presence of salicylate and absence of MarA and Rob suggests that other regulators like SoxS might also be able to play a role in regulating MIC. In fact, closely related species like *Salmonella* are known to have another regulator (homologous to MarA), RamA, which works together with *mar/sox/rob* towards conferring resistance [53]. Existence of such an additional regulator in *E. coli* is a likely possibility.

This study confirms the role of salicylate in altering the MIC of all three studied antibiotics via MarA/Rob respectively. We also studied if absence of either of MarA or Rob has the same effect on cells as absence of both. We observed that MIC in absence of either of MarA and Rob was same as that of wild-type cells (both in presence and absence of inducers) suggesting in absence of one, the other compensates for the loss of one of the regulators. Such redundancy in genetic network is quite ubiquitous and it is not surprising that it exists for a role as critical as stress response in bacterium *E. coli*.

### Cell density effect and quantification of cell death using Propidium Iodide (PI) assay

The previous section helped us understand the role of inducers in altering MIC. However, in absence of only one regulator (either MarA or Rob) it was difficult to capture any difference in MIC as the inducer salicylate acts via both MarA and Rob. In the double mutant, *ΔmarA Δrob* we notice maximum difference in MIC in presence as well as absence of salicylate. To further confirm the independent role of MarA and Rob, we quantified in response to exposure to antibiotics (Propidium Iodide fluoresces when in contact with DNA, thus is used to quantify fraction of cells which are dead). This will help us in understanding cell density effect in higher antibiotic exposure and also help quantify the independent roles of MarA and Rob.

Cells at higher densities (around 10^8^ cells per ml) were exposed to antibiotics in presence and absence of salicylate to track the percentage of dead cells using propidium iodide assay. We observed that in presence of salicylate, wild-type cells survive lethal antibiotics dosage for all three antibiotics studied (Figure 2, 3, and 4). As shown in Figure 2–4, the double mutant *ΔmarA Δrob* cells are susceptible to antibiotics even in presence of salicylate. In presence of carbenecillin antibiotic, wild-type cell showed survival in presence of salicylate till 40μg/ml of carbenecillin (Figure 2). *ΔmarA* cells were also able to survive exposure of antibiotics with the help of salicylate. However, *ΔmarA Δrob* cells showed ~50% percent dead cells, when exposed to 10 – 40μg/ml of carbenecillin in both absence and presence of salicylate. *Δrob* cells also showed similar behavior as *ΔmarA Δrob*, however, percentage of dead cells was lower (20–30%) as compared to the double mutant. Similar results were obtained for antibiotics cefotaxime and ciprofloxacin (Figure 3 and 4). When exposed to 2μg/ml cefotaxime, the percentage of dead cells in absence of MarA/Rob or MarA and Rob was 20–30%. However, at lower concentration of cefotaxime, presence of Rob helped in cell survival in absence of MarA (Figure 3) (cell death >10%). In case of ciprofloxacin, absence of MarA/Rob show similar effect, however, absence of both together renders cell susceptible to 1μg/ml ciprofloxacin similar to uninduced condition. Our result suggests that Rob plays an important role (more than that of MarA) in helping the cells survive antibiotic exposure.

**Figure 2.**
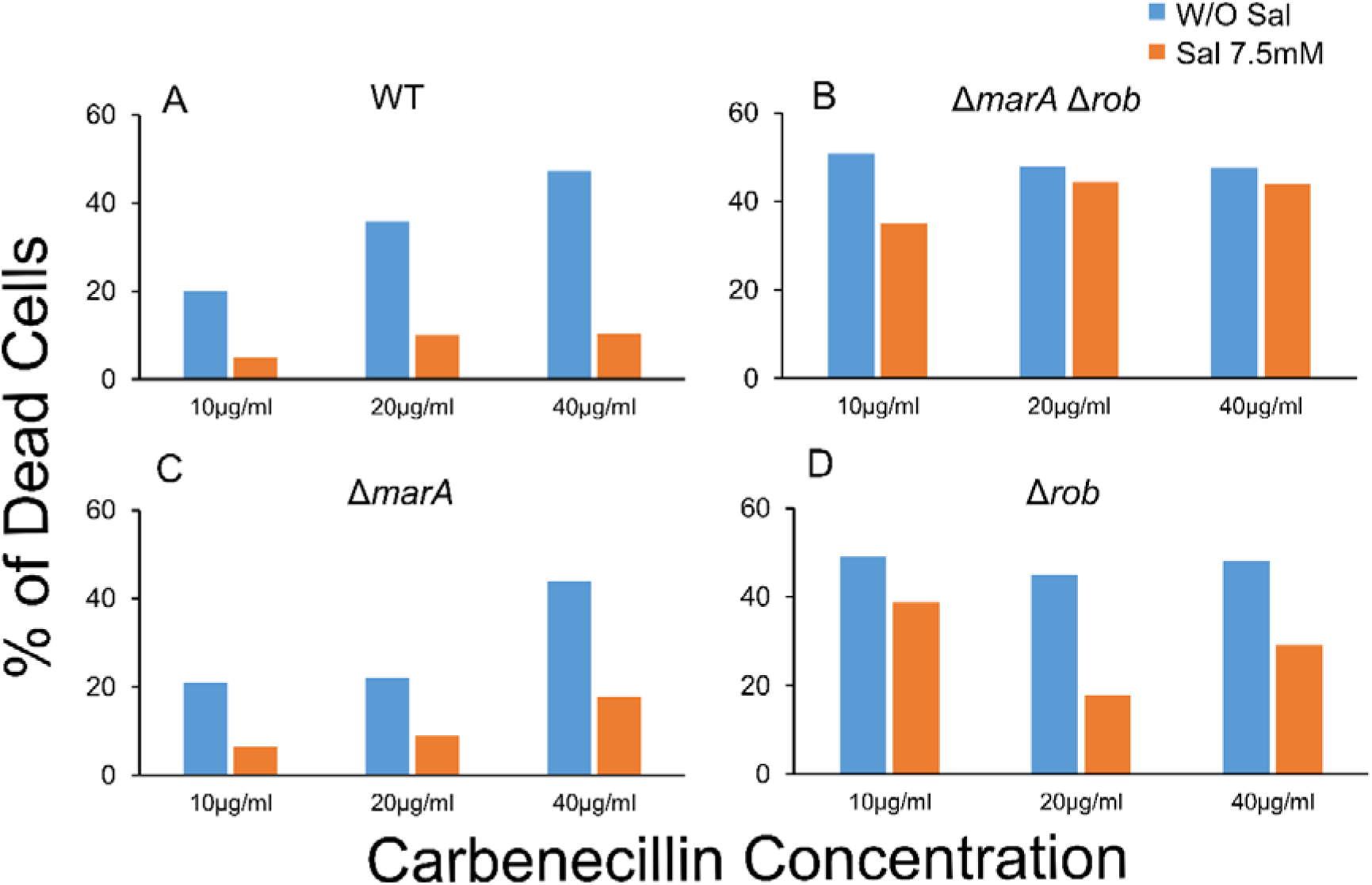
Percentage of dead cells in wild-type ***E. coli*** and mutants in presence of varying concentration of carbenecillin, with and without salicylate. (**A**) wild type (**B**) *ΔmarA Δrob* (**C**) *ΔmarA* (**D**) *Δrob*. Salicylate concentration used is 7.5mM and carbenecillin concentration used is 10μg/ml, 20μg/ml, and 40μg/ml.

**Figure 3.**
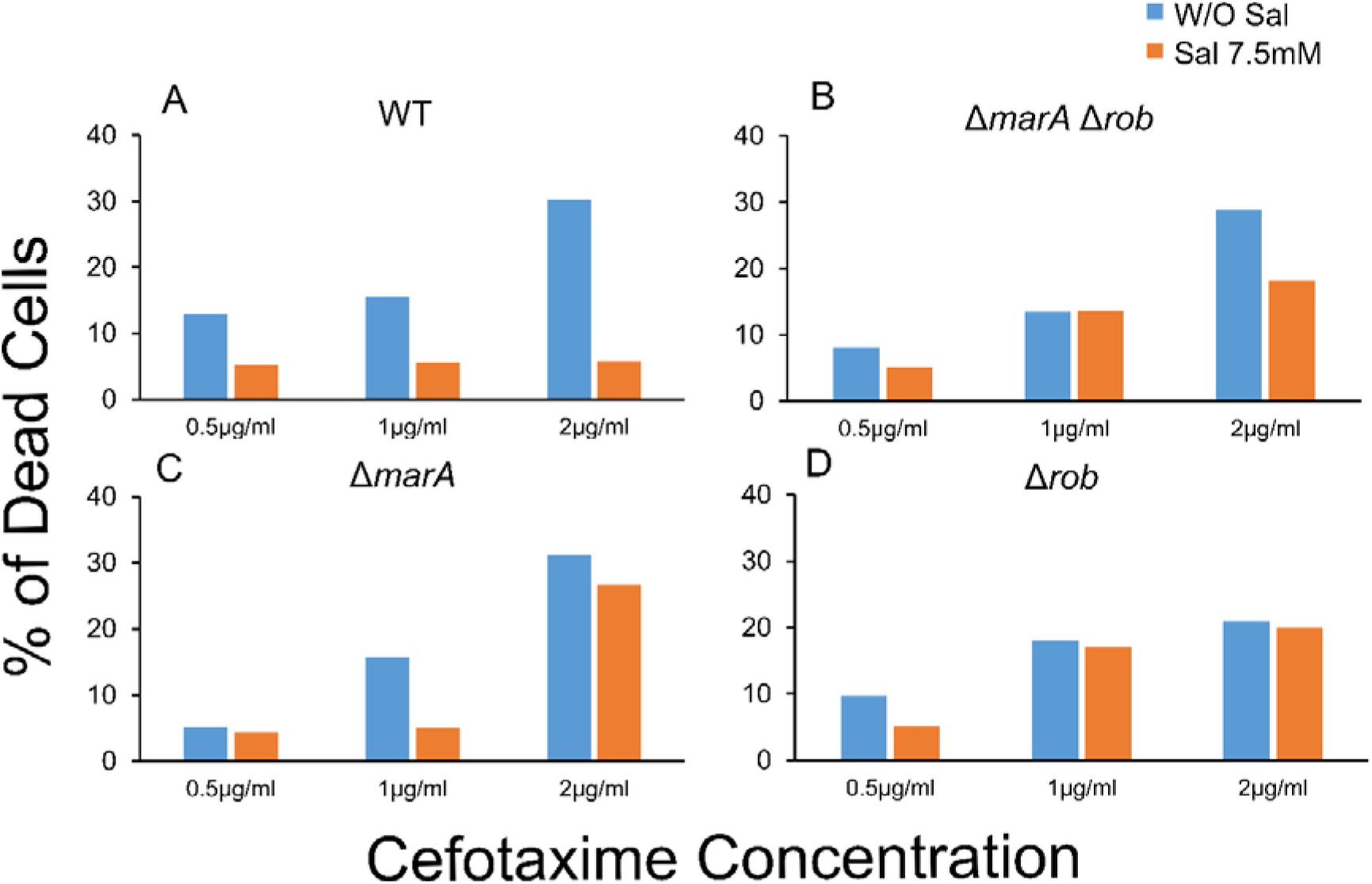
Percentage of dead cells in wild-type ***E. coli*** and mutants in presence of varying concentration of cefotaxime, with and without salicylate. (**A**) Wild type (**B**) *ΔmarA Δrob* (**C**) *ΔmarA* (**D**) *Δrob*. Salicylate concentration used is 7.5mM and cefotaxime concentration used is 0.5μg/ml, 1μg/ml, and 2μg/ml.

**Figure 4.**
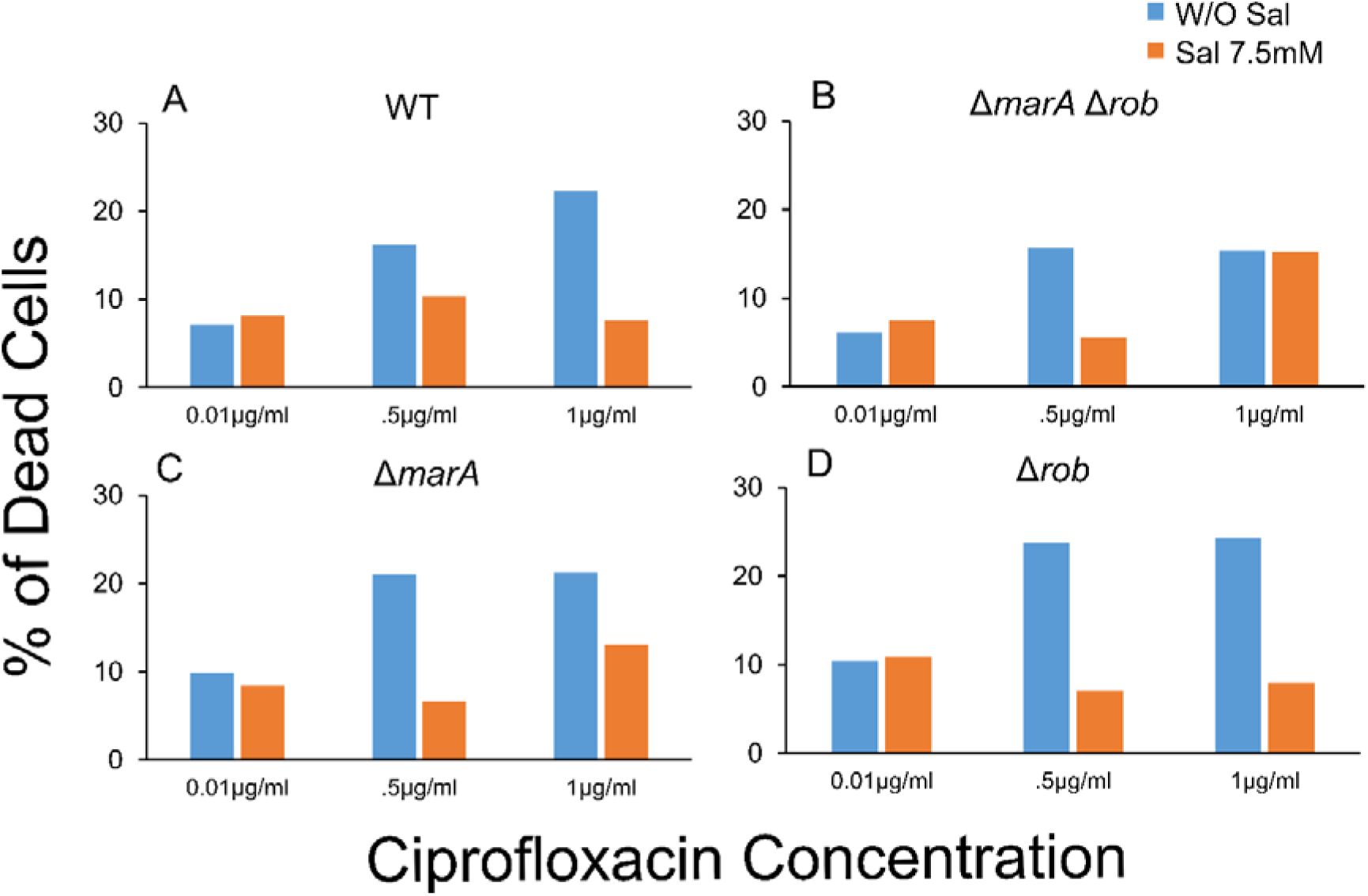
Percentage of dead cells in wild-type ***E. coli*** and mutants in presence of varying concentration of ciprofloxacin, with and without salicylate. (**A**) Wild type (**B**) *ΔmarA Δrob* (**C**) *ΔmarA* (**D**) *Δrob*. Salicylate concentration used is 7.5mM and ciprofloxacin concentration used is 0.01μg/ml, 0.5μg/ml, and 1μg/ml.

## Discussion

Our results suggest that Rob, like MarA, is an important player in responding to inducer salicylate. We quantify the phenotype associated with MarA and Rob in three different ways – growth kinetics, MIC, and PI assay. Growth experiments suggest that these systems are associated with a strong growth phenotype. MICs assays highlight the importance of chemicals like salicylate in altering the MICs of wide variety of clinically relevant antibiotics. PI assays help us understand the role of specific regulator in helping cells survive in an inducer dependent manner. MarA and Rob seems to act in coordination to help the cells survive wide range antibiotic stress in presence of salicylate. Rob being constitutively present in cell may enable cell to launch immediate response to harsh environment. The different mode of regulation of Rob as compared to MarA might be crucial to cell survival. Role of salicylate in independently acting via Rob can be of clinical importance as salicylate is the active component of medicines like aspirin.

Rob is reported to be present in 3–4 loci in cell with number varying from 5000–1000 molecules per cell [25,54]. In a single cell study, however, the number of Rob molecules in a cell was observed to be very low [55]. The variation in the number of protein molecules per cell suggest there could be cell-cell variation. Unlike other members of Arac/XylS regulators, the Prob is known to be regulated at post translational level [12,13]. It is present as agglomerate and dispersed to single molecule in presence of bipyridyl salt [34]. The transcriptional regulation of *Prob* in presence of salicylate is speculated but molecular mechanism of the same is not understood [35,36]. Moreover, the mechanism associated with transcriptional regulation of *Prob* by MarA, if any, is not understood [56,57]. Role of Rob protein, along with MarA and SoxS, in regulating cellular targets has been reported [7,12,13,22–24,32]. It is also speculated to be expressed in stationary phase under glucose and phosphate starvation [25,26]. What is the physiological relevance of Rob remains to be understood. It was first reported as right origin binding protein [25]. However, till date its role, if any, in DNA replication is not established. Our work attempts to understand role of Rob in cellular physiology, when exposed to wide variety of antibiotic stress. Further exploration of the mechanism involved in Rob mediated control of cellular physiology, in presence of inducers, can give new insights about intrinsic resistance mechanism.

